# Apusomonad photophobic behavior highlights cytoskeletal responses to blue light in early eukaryotes

**DOI:** 10.1101/2024.09.02.610796

**Authors:** Aika Shibata, Chiaki Yamamoto, Ryuji Yanase, Kogiku Shiba, Yu Sato, Akinori Yabuki, Kazuo Inaba

## Abstract

Light response is a fundamental characteristic of eukaryotic organisms, both unicellular and multicellular. However, no photoresponse has previously been reported in Apusomonadida, a group of small, free-living biflagellates phylogenetically positioned as a sister group to Opisthokonta (animals, fungi, and their unicellular relatives). Apusomonads are thus crucial for understanding the evolution of opisthokonts. Here, we report for the first time an avoidance response to blue light in the apusomonad *Podomonas kaiyoae*. This avoidance response is accompanied by an increase in gliding velocity, transient changes in flagellar waveforms, and alterations in cell shape. Dynamic cell contraction is induced by an increase in intracellular free Ca2+ and is inhibited by either a dynein inhibitor or an actin-disrupting drug. These findings suggest that the photophobic behavior of *Podomonas kaiyoae* relies on cytoskeletal responses mediated by both the dynein/tubulin and myosin/actin systems, which were acquired early in eukaryotic evolution. The dominance of the posterior flagellum in cilia-driven directional changes further supports the phylogenetic placement of apusomonads as a sister group to opisthokonts rather than other eukaryotic lineages.

## Introduction

In both aquatic and terrestrial environments, the behavior of many organisms is highly dependent on light. The growth and development of plants are governed by cellular processes controlled by photosynthesis, which involves sophisticated systems that respond to light (Franklin et al., 2005). In the ancestral Viridiplantae, the unicellular green alga *Chlamydomonas* exhibits positive and negative phototaxis as well as photoshock responses, mediated by a light-induced depolarization cascade initiated by channelrhodopsins. This cascade alters the waveforms or beating balance of its two morphologically identical flagella (Hegemann and Berthold, 2009). In animals, the visual system responds to environmental light through sensory organs rich in type-II rhodopsin photoreceptors, transforming light information into movement via the motor neuron system (Jékely, 2009).

The mechanisms of photoreception had already been acquired in the early stage of eukaryotic evolution. Excavata is a diverse collection of ancestral unicellular eukaryotes, such as Discoba and Metamonada (Simpson et al, 2017; Jewari et al, 2023). *Euglena*, a member of Discoba, exhibits a unique light-responsive movement known as euglenoid movement (Murray, 1981; Suzaki and Williamson, 1985). This movement involves both the Ca^2+^-dependent change in the direction of flagellar propagation and a large deformation of the cell, accompanied by contraction of a flexible, protein-based structure called the pellicle. Another major eukaryotic group, the unikonts – recently termed “Amorphea” (Burki et al., 2020) – includes Amoebozoa and Obazoa (Simpson et al., 2017). This is a group that consists of Amoebozoa and Obazoa (Simpson et al, 2017). The latter lineage contains fungi and animals (metazoans), which have evolved a multicellular visual organ connected to the central nervous system (Jékely, 2009). Early diverging fungal representatives, such as chytrids, consist of several phyla of unicellular fungi that produce posteriorly flagellated motile spores. Some chytrid species show clear positive phototaxis (Robertson, 1972; Avelar et al., 2014; Swafford and Oakley, 2018). Light reception in these organisms is mediated by an eyespot-like, lipid-rich structure known as the side-body complex or the microbody-lipid globule complex, which regulates flagellar beating (Avelar et al., 2014; Galindo et al., 2022). Phototaxis in amoebozoans has been observed in the amoeboid cells and plasmodia of *Dictyostelium* and *Physarum* (Francis, 1964; Rakoczy, 1973), but has not yet been demonstrated in the flagellated cells of *Physarum*. These observations suggest that light-induced changes in cell motility in early eukaryotes likely involved coordination between ciliary or flagellar movement and dynamic changes in cell shape mediated by microtubule- or actin-filament-based motility (Jékely, 2009; Fritz-Laylin, 2020).

Apusomonads are heterotrophic flagellates that occur both in marine and terrestrial habitats. They locomote mainly by gliding on surfaces which mostly depends on movement by the posterior flagellum together with the pseudopodia, the beating of its anterior flagellum does not appear to be involved in the movement (Cavalier-Smith and Chao, 2010; Heiss et al, 2013; 2017). Recent phylogenomic studies suggest that apusomonads are the sister group to opisthokonts and both are positioned within the group Obazoa together with Breviata (Brown et al, 2013). Despite the phylogenetic position and motility characteristics of apusomonads, which suggest they retain ancestral eukaryotic features, no phototactic behavior or photoreceptors have been described in these organisms. This lack of information represents a significant gap in our understanding of the cytoskeletal roles in photoresponse during eukaryotic evolution, particularly in a group that links Amoebozoa and Obazoa (Jékely, 2009). In this study, we present the first report of a photoresponse in an apusomonad in the recently identified species *Podomonas kaiyoae* (Yabuki et al, 2023). We explore the cytoskeletal dynamics involved in the photoresponse of this representative species from an early-branching eukaryotic lineage.

## Results

### Photophobic response of the apusomonad *Podomonas kaiyoae*

To determine if apusomonads display any light-responsive behavior, *Podomonas kaiyoae* cells were observed under stroboscopic red LED light, followed by exposure to blue light at the center of the field. The behavior of the cells was recorded under a phase contrast microscope. Upon blue-light illumination, the distribution of *P. kaiyoae* cells decreased gradually within the illuminated area, creating a distinct empty circle in the center after 100 seconds (Fig. 1a, Supplementary Video 1). Most cells in the illuminated area exhibited a photophobic response characterized by cell contraction and extension. When the blue light was turned off, the cells gradually reoccupied the previously empty area, indicating that the reduction in cell numbers within the center field was due to an avoidance response (Fig. 1a, Supplementary Video 1). The avoidance response was most effectively triggered by blue light (440 nm) but could also be induced by cyan light (470 nm) and, to a lesser extent, by green light (550 nm) (Fig. 1a bottom; Fig. 1b, 1c; Supplementary Video 2).

**Fig 1.**
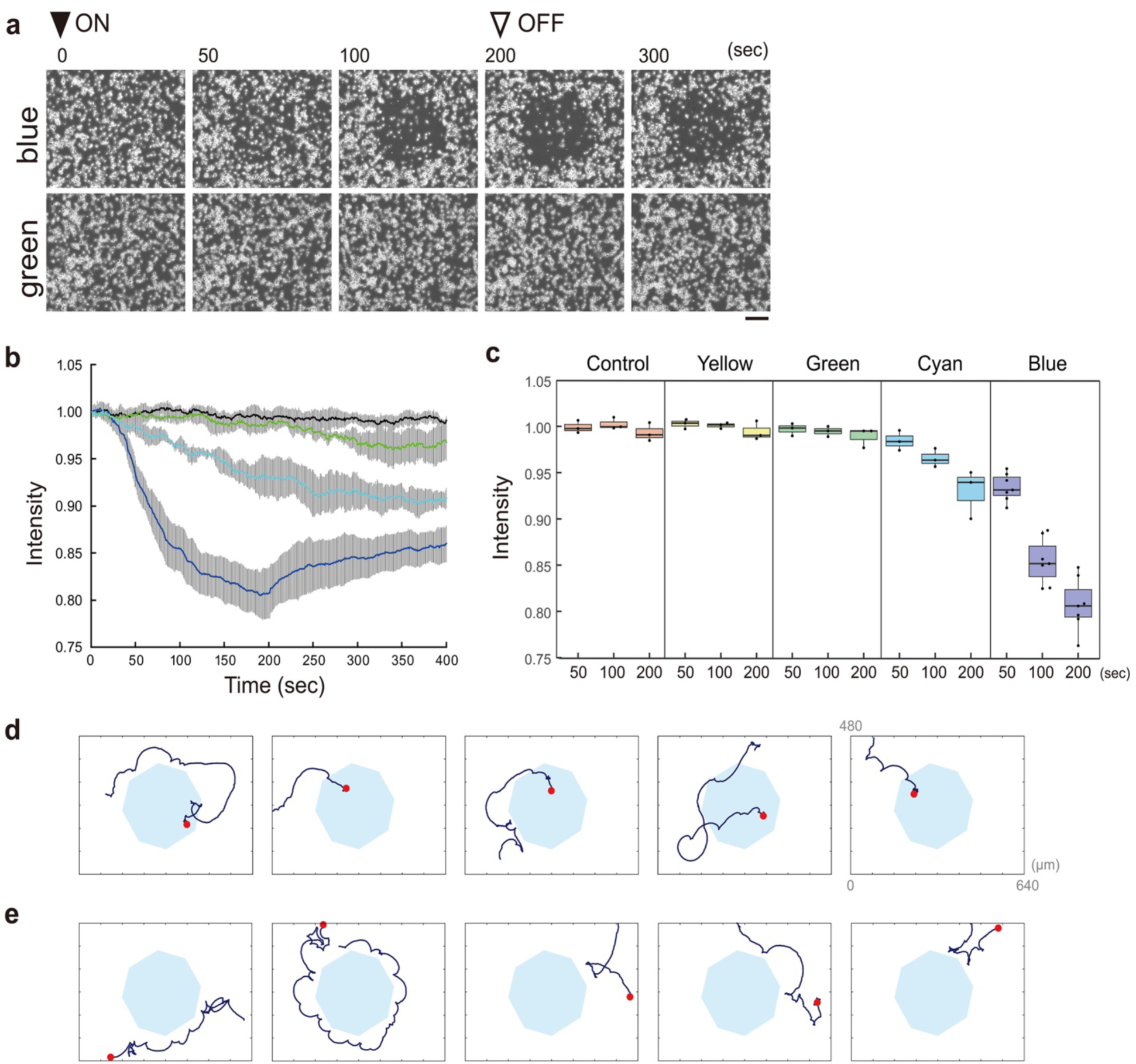
Light-induced avoidance response of the apusomonad *Podomonas kaiyoae*. **(a)** Sequential images show *P. kaiyoae* cells avoiding an illuminated field. The images capture the moments when the blue or green light is turned on (ON) and off (OFF). The scale bar represents 100 µm. **(b)** Graph displaying changes in image intensity at the center of the field under different light colors. Light was turned on at 0 seconds and off at 200 seconds. The colors of the curves correspond to the colors of the lights used, with the black curve representing the control (no illumination). Error bars represent standard deviation, with a sample size of N=3. **(c)** Effect of various light colors on the decrease in intensity within the illuminated field. **(d)** Behavior of a cell gliding from a bright to a dark field across the blue light border. The cell does not exhibit a step-down response. **(e)** Behavior of a cell gliding from a dark to a bright field across the blue light border, showing a clear step-up response. In both (d) and (e), cell trajectories were recorded for 34-100 seconds. The long side of the rectangle in the images measures 640 µm, and the red spot marks the initial position.

The response was observed in a light-intensity-dependent manner (Fig. 1c). Cells passed through the boundary between bright and dark areas as they escaped from the illuminated region (Fig. 1d, Supplementary Video 3). However, when re-entering the illuminated area, cells often stopped gliding at the edge of the light and changed direction away from the illuminated region (Fig. 1e, Supplementary Video 4). These findings suggest that the response to light is a step-up photophobic response. A typical avoidance response pattern under bright field microscopy included a sequential process: cessation of gliding, a ∼90-degree turn, and resumption of gliding until the cell crossed the bright-dark boundary.

To examine whether *P. kaiyoae* cells show positive phototaxis, blue light was illuminated from one side of a chamber on a glass slide including cells. Although the cells became scarce near the light source under blue light illumination, we could not observe any clear phototactic movement toward the light source (Supplementary Figure 1 and Supplementary Video 5).

### Cell contraction and changes in gliding direction are necessary for photophobic response

Two types of *P. kaiyoae* cells were observed in terms of motility. Before illumination under red light, many cells were settled on the glass surface using their posterior pseudopodia and occasionally exhibited contraction-extension movements (termed ‘settled’). Only a small population of cells displayed progressive gliding motility (termed ‘gliding’) with occasional changes in direction due to cell contraction-extension, as previously reported in *P. kaiyoae* and other apusomonads (Yabuki et al., 2023; Cavalier-Smith and Chao, 2010; Heiss et al., 2013). Upon blue light illumination, many settled cells detached and began gliding (Fig. 2a, top; Supplementary Video 6). During blue light illumination, gliding cells frequently changed direction with cell contraction (Fig. 2a, bottom; Supplementary Video 7).

**Fig 2.**
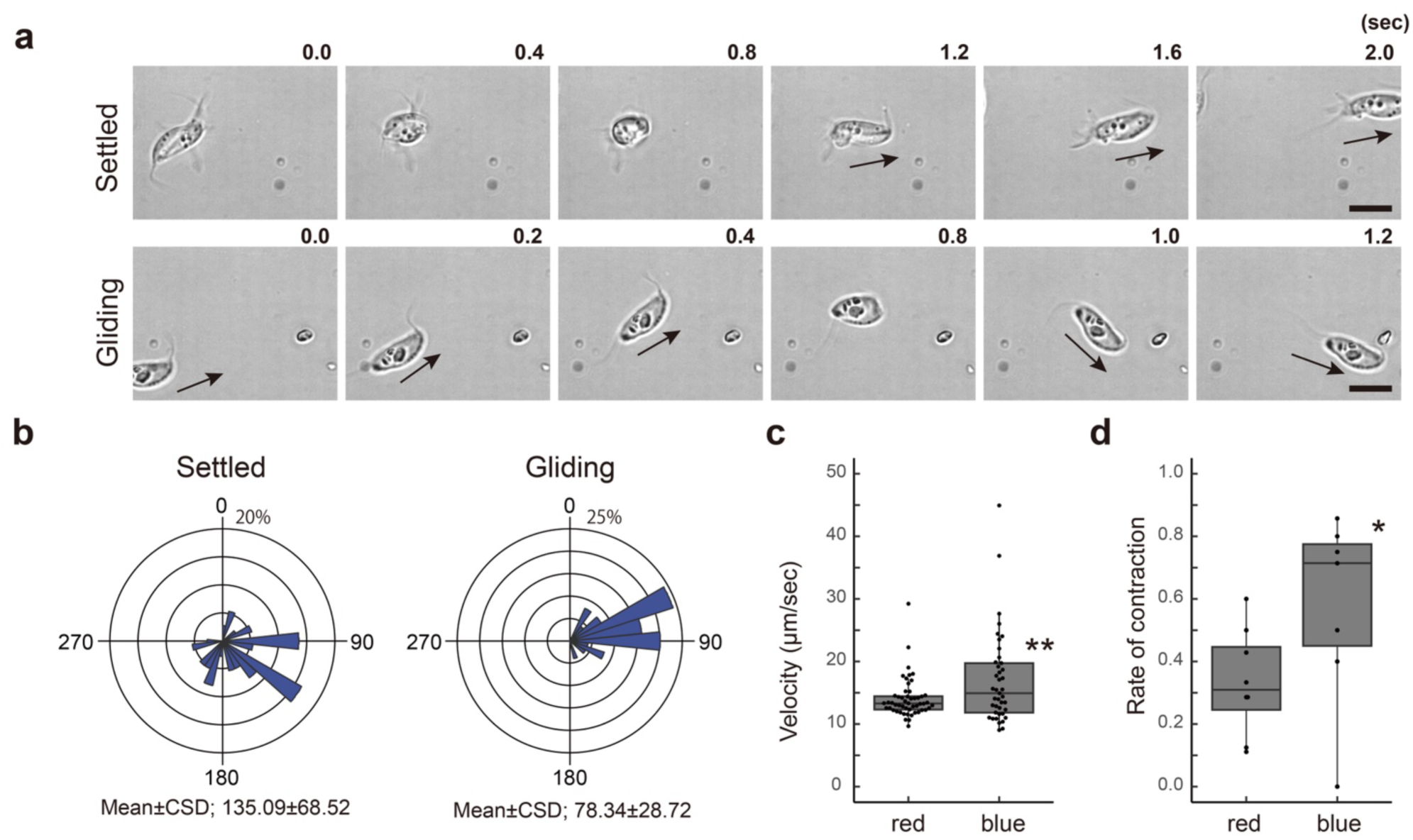
Light-induced activation of gliding and cell contraction in apusomonads. **(a)** Both settled cells (top) and gliding cells (bottom) on a glass surface contract in response to blue light illumination. Arrows indicate the direction of gliding. The scale bar represents 10 µm. **(b)** Rose plots depict the angle displacement in gliding direction post-contraction under blue light illumination. The left plot represents cells that settled and started gliding after contraction, with the angle displacement measured between the anterior-posterior axis before contraction and the gliding direction post-contraction (n=37). The right plot represents cells that were already gliding and altered their direction post-contraction (n=25). Circular standard deviation (CSD) is noted. **(c, d)** Box plots illustrate gliding velocity (c) and contraction rate (d) under red or blue light. ** Significant at p < 0.01, * p < 0.05 (Student’s t-test) when compared to red light.

To test if blue-light-dependent cell contraction and the initiation of gliding are correlated with directional blue-light avoidance movement, changes in cell orientation before and after blue light illumination were measured. The rose plots show that settled cells began gliding in various directions relative to their orientation before illumination (Fig. 2b, left). However, in gliding cells, the direction became limited to 60-90° relative to their pre-illumination orientation (Fig. 2b, right). The circular standard deviation demonstrated that the gliding direction was random in settled cells but significantly oriented in gliding cells (Fig. 2b). Blue light illumination activated the gliding movement of *P. kaiyoae* cells, increasing their gliding velocity to approximately 1.2 times the pre-illumination speed (Fig. 2c). The frequency of contraction also increased to about 1.7 times the total cell population after blue light illumination (Fig. 2d).

### The posterior flagellum is linked to cell contraction

The photophobic response in *P. kaiyoae* involves changes in cell shape, including contraction and subsequent extension for re-gliding. To understand the cellular mechanism underlying blue-light-induced contraction and gliding, the dynamics of microtubules and actin filaments were examined. Immunofluorescent observation using an anti-acetylated α-tubulin antibody revealed a microtubule network that included the anterior and posterior flagella and the dorsal roots (Fig. 3a, Supplementary Figure 2). The anterior flagellum continuously beats in a three-dimensional fashion (Supplementary Video 6 and 7). Staining with Alexa-labeled phalloidin showed actin filaments in both the anterior cytoplasm and trailing pseudopodia. The trailing pseudopodia extended several filamentous projections that adhered to the glass surface. The lengths and numbers of pseudopodia varied among cells (Supplementary Figure 2).

**Fig 3.**
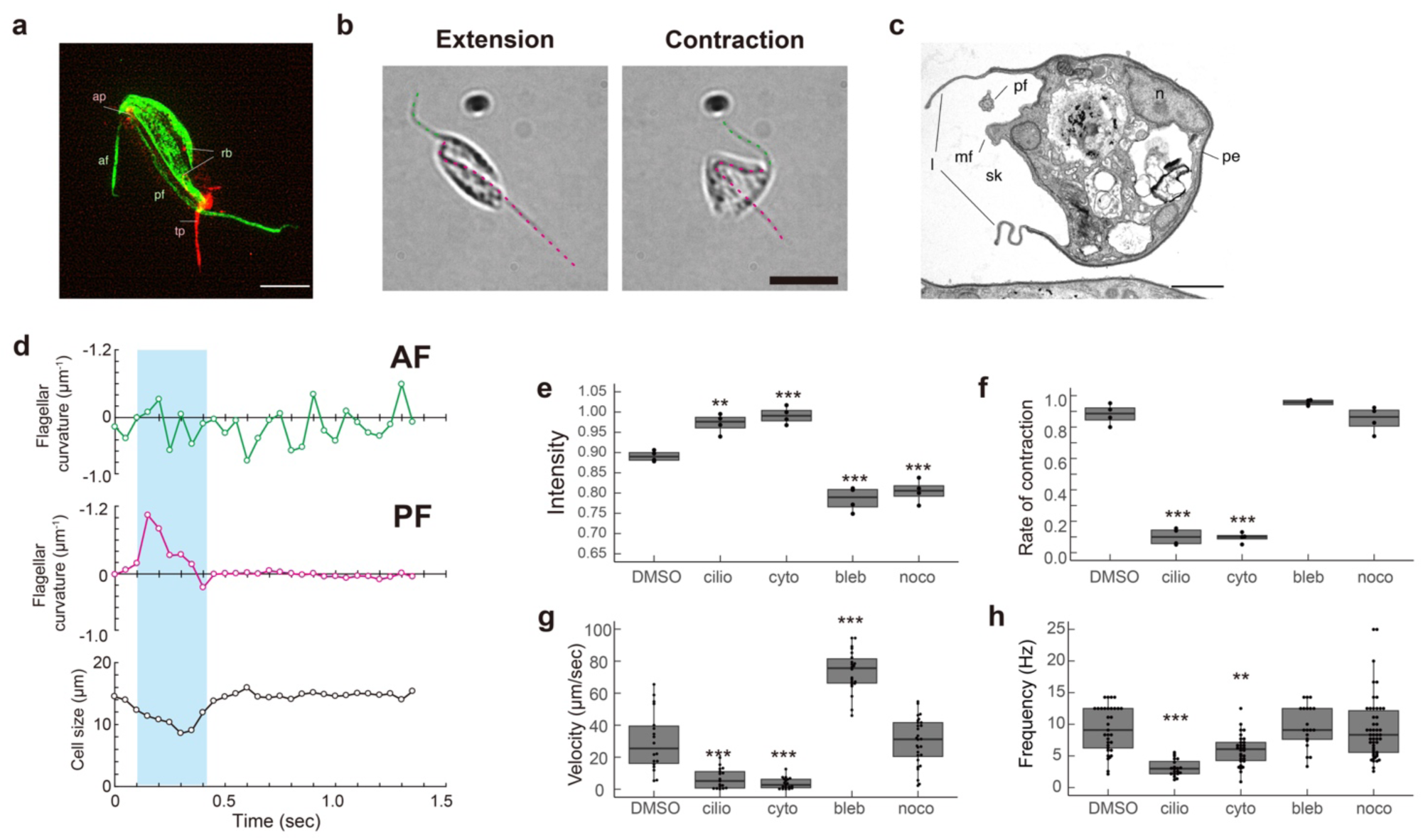
Roles of cytoskeletal structures in the avoidance response of apusomonads. (a) Cytoskeletal images of *P. kaiyoae* stained with anti-acetylated tubulin (green) and phalloidin (red). Key structures include the anterior flagellum (af), posterior flagellum (pf), ribbon (rb), anterior pseudopodium (ap), and trailing pseudopodium (tp). The scale bar represents 5 µm. (b) Two morphological states of *P. kaiyoae* are depicted: extension (left) and contraction (right). The anterior and posterior flagella are marked with dashed lines in green and magenta, respectively. The scale bar represents 10 µm. (c) Thin-sectioned electron micrograph of *P. kaiyoae*, showing the posterior flagellum ventrally positioned along the cell body within a fold called the skirt, formed by two lips. Other structures include the posterior flagellum (pf), lip (l), skirt (sk), microfilament bundle (mf), nucleus (n), and pellicle (pe). The scale bar represents 1 µm. (d) Changes in the maximum curvature of the anterior and posterior flagella, measured 2.5 µm from the flagellar base during cell contraction. Cell size is expressed as the distance between the anterior and posterior ends of the cell body. (e-h) Effects of cytoskeletal and motor-related drugs on various aspects of cell behavior. (e) Intensity measured as brightness at the center area under illumination. (f) Contraction rate per cell. (g) Gliding velocity 5 seconds post-illumination. (h) Beat frequency of the anterior flagellum. Distribution of values is plotted in a box plot. *** Significant at p < 0.001, ** p < 0.01 (Dunnet test) when compared with DMSO.

The posterior flagellum lay through the center of the cell in the extended state (Fig. 3b, left), with the portion protruding from the cell body slightly bent in both contraction and extension states (Fig. 3b; Supplementary Video 8). During the contraction state, the flagellum bent significantly towards the ventral side of the cell (Fig. 3b, right; Supplementary Video 8). In some cases, the posterior flagellum bent significantly and extended out from the cell body, indicating that the bending of the posterior flagellum was linked to contraction.

Thin-sectioned electron microscopy showed that most of the dorsal portion was covered by an electron-dense pellicle. The anterior flagellum protruded through a sleeve posteriorly connected to a fold termed a skirt. The posterior flagellum was positioned ventrally along the cell body, engulfed by the lips of the skirt (Fig. 3c, Supplementary Figure 3). The protruded ventral surface featured a row of cortical singlet microtubules (ribbon) aligned in the anterior part (Supplementary Figure 3). No distinct projections, including dyneins, were observed on these singlet microtubules. This area also appeared protruded in the posterior part, where microfilaments were present (Fig. 3c, Supplementary Figure 3). The cytoskeletal architecture was not identical but similar to that of *Thecamonas trahens* (Heiss et al., 2013).

To correlate flagellar motility with the mechanism of cell contraction, the bending of anterior and posterior flagella during cell contraction was measured (Fig. 3d). Results show that during cell contraction-extension, the anterior flagellum continuously beats with curvature oscillation, while the posterior flagellum remains almost straight with less bending during the extension state. However, the curvature transiently increased during cell contraction (Fig. 3d).

### Cooperation of two cytoskeletal systems in contraction-extension transition

To understand the roles of cytoskeletal dynamics in the contraction-extension response in *P. kaiyoae*, the effects of drugs targeting the cytoskeleton and its molecular motors were examined (Fig. 3e-h). The microtubule polymerization inhibitor nocodazole was used first. Treatment with nocodazole at 12.5 µM for 15 minutes did not significantly inhibit the rate of contraction (Fig. 3f), gliding velocity (Fig. 3g), or anterior flagellar beat frequency (Fig. 3h). The photoavoidance response was only slightly activated under nocodazole treatment (Fig. 3e; Supplementary Video 9).

Cells were also treated with cytochalasin B, an actin polymerization inhibitor, which inhibited the photoavoidance response after 15 minutes of incubation at 200 µM, with no decrease in flagellar beat frequency (Fig. 3e and 3h, Supplementary Video 10). However, cytochalasin B significantly inhibited both the rate of contraction and the gliding velocity (Fig. 3f and 3g).

Inhibitors for molecular motors were also tested. Ciliobrevin, a dynein inhibitor (Firestone et al., 2012), significantly inhibited both the photoavoidance response and the rate of contraction at 40 µM (Fig. 3e and 3f; Supplementary Video 11), similarly to cytochalasin B treatment. As expected, the flagellar beat frequency was greatly reduced (Fig. 3h). Similar to cytochalasin B, ciliobrevin treatment significantly lowered the gliding velocity (Fig. 3g). Lastly, cells were treated with blebbistatin, a pharmacological compound that blocks myosin II activity (Straight et al., 2003). Blebbistatin had no effect on cell contraction or anterior flagellar beat frequency (Fig. 3f and 3h). The photoavoidance response was slightly activated by blebbistatin (Fig. 3e), but notably, the compound significantly increased cell gliding velocity (Fig. 3g, Supplementary Video 12).

Under red light, many cells settled on the glass surface using their posterior trailing pseudopodia. This settling behavior persisted in some cells even immediately after blue light illumination (Fig. 2a, top; Supplementary Video 6). Upon blue light exposure, the trailing pseudopodia of the cells detached and were drawn along by the cell body during gliding (Supplementary Video 6). Trailing pseudopodia were observed in cells treated with nocodazole or blebbistatin (Supplementary Video 9 and Video 12). However, in cells treated with cytochalasin or ciliobrevin, the pseudopodia retracted into the cell body (Supplementary Video 10 and Video 11). Furthermore, these cells exhibited reduced adhesion to the glass surface, with some cells flipping over (Supplementary Video 13 and Video 14). Cell flipping occurred more frequently in ciliobrevin-treated cells than in cytochalasin-treated ones. In the latter case, the posterior flagellum often remained attached to the glass slide, preventing flipping (Supplementary Video 13).

### Ca^2+^ is essential for both progressive gliding and cell contraction

Calcium ions (Ca²⁺) are crucial signaling molecules for regulating cellular phototaxis and chemotaxis (Cai and Clapham, 2012; Inaba, 2003; 2015). To investigate the regulatory mechanisms underlying the photophobic response in *P. kaiyoae*, we examined the effects of Ca²⁺ on forward gliding movement and cell contraction. Initially, cells were transferred into Ca²⁺-free (EGTA-chelated) artificial seawater medium, which resulted in significant cellular damage and disruption. Consequently, we employed a low Ca²⁺ artificial seawater medium containing one-fourth the normal Ca²⁺ concentration (2.5 mM). Under this low Ca²⁺ condition, cells exhibited a dramatic loss of forward motility, cell contraction, and gliding (Fig. 4a–d). Instead, *P. kaiyoae* cells under low Ca²⁺ displayed a pivot-like movement, similar to the motility patterns observed in ciliobrevin- or cytochalasin-treated cells (Supplementary Video 15). Both anterior and posterior flagella appeared to be beating, but at a significantly reduced frequency (Supplementary Video 16). Forward gliding motility and cell contraction were restored when the cells were transferred back to artificial seawater (ASW) (Fig. 4a–d).

**Fig 4.**
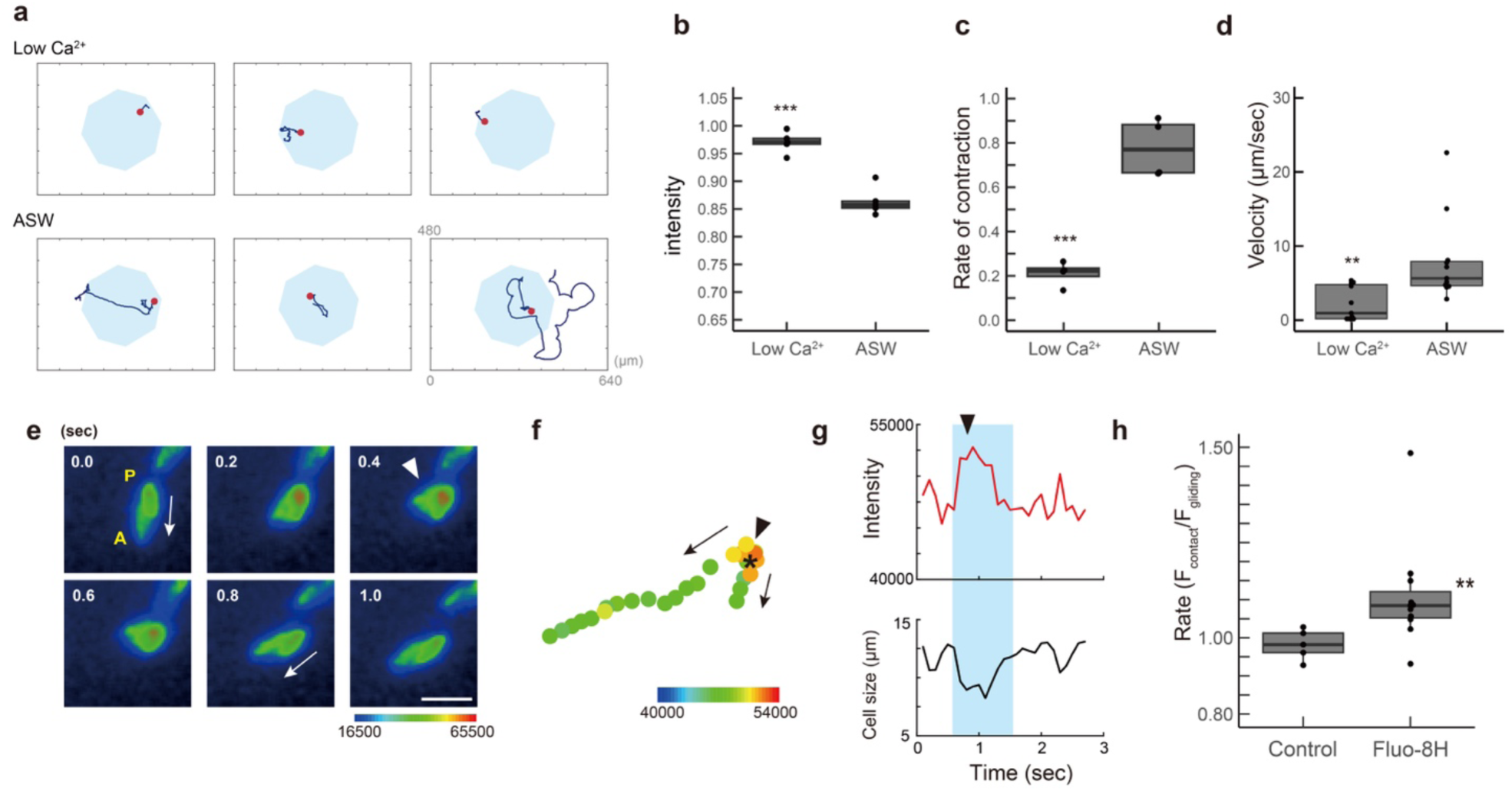
Roles of Ca^2+^ in the regulation of the avoidance response in apusomonads. (a) Cells were first observed in a low Ca^2+^ ASW. Then the solution was substituted back to ASW. Cell trajectories recorded for 87-100 seconds. The long side of the rectangle measures 640 µm, with the red spot indicating the initial position. **(b-d)** Calcium (Ca²⁺) is crucial for cell contraction and gliding. (b) Intensity measured as brightness at the center under illumination. (c) Contraction rate per cell. (d) Gliding velocity 5 seconds post-illumination. These variables are plotted as box plots, comparing conditions in low Ca²⁺ ASW (Low Ca) versus standard ASW. ** Significant at p < 0.01, * p < 0.05 (Student’s t-test) when compared with the value in ASW. (e) Changes in intracellular Ca²⁺ concentration ([Ca²⁺]_i_) during cell contraction. Fluo-8H fluorescence is displayed using a pseudocolor lookup table (LUT). Arrows indicate gliding, and arrowheads indicate contractions. The anterior (A) and posterior (P) ends of the cell are marked. The scale bar represents 10 µm. (f) Cell trajectories and maximum [Ca²⁺]_i_ intensity signals along the cell axis are displayed in pseudocolors. Arrows and arrowheads indicate gliding and contractions, respectively. An asterisk marks the initial position. (g) Changes in maximum fluorescence intensity during cell contraction, shown in light blue. An increase in [Ca²⁺]_i_ is observed during contraction, with the arrowhead indicating the same position as in (e) and (f). (h) Rate of [Ca²⁺]_i_ increase during cell contraction in Fluo-8H unloaded (Control) versus loaded cells. F_contract_ and F_gliding_ represent maximum [Ca²⁺]_i_ intensity signals along the cell axis at minimum and maximum cell lengths, respectively. ** Significant at p < 0.01 (Student’s t-test) when compared with the Control.

To visualize Ca²⁺ dynamics during the photophobic response, we introduced a Ca²⁺-selective fluorescent indicator, Fluo-8H, into *P. kaiyoae* cells. A membrane-permeable acetoxymethyl ester form of Fluo-8H was loaded into the cells in the presence of the non-ionic detergent Cremophor. Fluorescence emission was measured under cyan light (470 nm) excitation, which simultaneously induced the photophobic response. We observed that the fluorescence intensity of a cell increased a few seconds after cyan light irradiation (Fig. 4e and 4f, Supplementary Video 17), in parallel with cell contraction (Fig. 4g). The fluorescence intensity was not uniform across the cell; instead, a specific area, likely the posterior region, exhibited an increase in intensity. When the cell resumed forward gliding, the fluorescence returned to its initial levels. To exclude the possibility that changes in cell thickness affected the fluorescence intensity, we compared the ratio of maximum fluorescence intensity during gliding and contraction states between Fluo-8H-loaded cells and unloaded cells (autofluorescence only). The ratio was significantly higher in Fluo-8H-loaded cells (Fig. 4h), indicating that the observed changes in fluorescence intensity were due to intracellular Ca²⁺ concentration.

## Discussion

We describe for the first time a clear photoresponse in an apusomonad species. *P. kaiyoae* was originally isolated from a deep-sea sediment (Yabuki et al., 2023). It is known that bioluminescence plays a variety of biological roles in the communication among deep-sea organisms (Haddock, 2010). *P. kaiyoae*, like other apusomonads, is a heterotrophic protist that shows no photosynthesis but feed on bacteria and other small protists (Yabuki et al., 2023). The avoidance response of *P. kaiyoae* is evidently thought not to be used for feeding luminescent bacteria, but rather for escaping from the predators, presumably large luminescent protists or deep-sea deposit feeder. Alternatively, they may escape from faint light reaching to the surface of deep seabed to hide. It is also possible to consider that *P. kaiyoae* is not only found in deep sediments but also in shallow waters. In this case, the photoavoidance response would help the cells to orientate themselves in the sediments, or to protect themselves from near UV or blue light.

While the activation of forward gliding by blue light may induce a decrease in the population of *P. kaiyoae* cells in the illuminated area, suggesting photokinesis, statistical analysis of gliding movement across the bright-dark boundary clearly indicates a photoavoidance behavior (Fig. 1a). Moreover, the directional change in gliding was more pronounced when cells moved from dark to bright areas, but less evident when moving from bright to dark areas, suggesting a step-up response in the photophobic behavior (Fig. 1d and 1e).

In this study, we did not identify any specific photoreceptors or light-sensing subcellular structures. Photoreception is a fundamental characteristic in both prokaryotes and eukaryotes, with photoreceptor proteins being the primary molecular players involved (Wan and Jékely, 2021). Photoreceptor proteins, including proteorhodopsin (type I) and various vertebrate (type II) opsins, are widely distributed across organisms. Unicellular eukaryotes are also known to possess several types of photoreceptor proteins and associated cellular structures (Jékely, 2009). For instance, zoospores of the chytrid fungus *Blastocladiella emersonii*, a close relative within opisthokonts, exhibit phototactic swimming behavior in response to green light, mediated by a unique fusion protein containing a type I rhodopsin (Saranak and Foster, 1997; Yu and Fischer, 2019; Galindo et al., 2022). In the apusomonad *Thecamonas trahens*, genomic analyses suggest the presence of potential photoreceptors and light-perception-related molecular machinery, including a cyclic nucleotide-gated channel (Galindo et al., 2022) and proteins with BLUF domains (photoactivated adenylate cyclase PAC) (Iseki et al., 2002; Kaushik et al., 2019) or LOV domains (PASK-PAS) (Cosson et al., 2003; Xing et al., 2023). The cyclic nucleotide-gated channel, which responds to rhodopsin-mediated cGMP production, controls flagellar beating in *B. emersonii* (Galindo et al., 2022). Our findings indicate that blue light is the most effective wavelength for inducing photoavoidance, suggesting the presence of blue light photoreceptors in *P. kaiyoae*.

Ancestral eukaryotes underwent significant evolutionary events, including the acquisition of novel cytoskeletal motility systems (Cavalier-Smith, 2002, 2006; Brown et al., 2013; Yubuki and Leander, 2013). This study provides important insights into the evolution of cytoskeletal roles in motility. The photoresponse in *P. kaiyoae* consists of two steps: activation of forward gliding and cell contraction, which induces a change in gliding direction. The microtubule-depolymerizing drug nocodazole did not inhibit gliding or cell contraction, whereas the dynein inhibitor ciliobrevin blocked both types of motility. A similar effect of a dynein inhibitor has been observed in trypanosomes (Broadhead et al., 2006; Gadelha et al., 2007; Sun et al., 2018). These findings suggest that the photophobic response is mediated by dynein-driven flagellar motility rather than by the dynamics of singlet microtubule structures, such as ribbon structures. Additionally, since ciliobrevin and cytochalasin blocked both types of motility, actin dynamics are also likely necessary for cell gliding and contraction. Retraction of trailing pseudopodia was observed in cells treated with both drugs, indicating that pseudopodia may serve as anchors during cell contraction and turning.

Drug-induced changes in cell behavior demonstrate that both flagella and actin filaments are responsible for gliding, suggesting a cooperative interaction between these two cytoskeletal systems during gliding and cell contraction. Observations of cytochalasin-treated cells (Supplementary Video 13) indicate that the posterior flagellum plays a role in attachment and possibly in cell gliding. It is known that flagellar surface-based gliding is driven by intraflagellar transport (IFT) in *Chlamydomonas* (Bloodgood, 1981; Shih et al., 2013), a movement also observed in sea urchin larvae cilia (Kamiya et al., 2018). Therefore, the posterior flagellum may contribute to cell contraction through large asymmetric bending and to gliding on the glass surface via IFT.

In contrast, amoeboid movement is a common mode of gliding in eukaryotes (Taylor and Condeelis, 1979). The anterior actin-rich pseudopods function as the leading edge in the amoeboid excavate *Naegleria* (Fritz-Laylin et al., 2010), unicellular slime mold (Swanson and Taylor, 1982; Ueda and Ogihara, 1994), and the zoospore of chytrid fungi (Fritz-Laylin et al., 2017). Thus, it is possible that the dynamics of the anterior actin network contribute to gliding in coordination with IFT in the posterior flagellum. Myosin may be involved not in generating the gliding force but rather in anchoring the posterior pseudopodia. Considering that blebbistatin significantly increases gliding velocity (Fig. 3g), myosin likely provides mechanical resistance during gliding in *P. kaiyoae*, contrasting with its role in neutrophils (Eddy et al., 2000).

In conclusion, *P. kaiyoae* exhibits a cooperative movement involving two ancestral motility systems: IFT-dependent ciliary surface motility and amoeboid movement based on actin dynamics. This motility system, observed in *P. kaiyoae*, may reflect the motility mechanisms used by the ancestor of opisthokonts, considering cytoskeleton-based motility in early opisthokonts, such as chytrid fungi (Fritz-Laylin et al., 2017; Prostak et al., 2021) and early metazoans like choanoflagellates (Brunet and King, 2017; Brunet et al., 2019).

The Ca²⁺-dependent flagellar asymmetry in *P. kaiyoae* is consistent with the properties observed in metazoans and fungi (Inaba, 2015), supporting the classification of Apusomonadida as the sister group of Opisthokonta (Cavalier-Smith and Chao, 2003; Adl et al., 2012; Brown et al., 2013). *P. kaiyoae* exhibits a large bend in the posterior flagellum at high Ca²⁺ concentrations, whereas *Trypanosoma* reverses the direction of wave propagation in the anterior flagellum to a tip-to-base pattern under high Ca²⁺ concentrations, with an asymmetric form (Holwill and McGregor, 1976). Animal sperm propagate asymmetric waves in the posterior flagellum at high intracellular Ca²⁺ concentrations (Inaba, 2015). A similar Ca²⁺-dependent asymmetric movement of the posterior flagellum was observed in the zoospore of the fungus *Blastocladiella emersonii* (Miles and Holwill, 1969), although the conversion of waveform symmetry was extremely slow. Our findings demonstrate that a large asymmetric bending of the posterior flagellum is induced by an increase in intracellular Ca²⁺ concentration. Genomic surveys of eukaryotes suggest that apusomonads have already acquired a set of molecules necessary for Ca²⁺ response, such as the sperm-specific Ca²⁺ channel CatSper (Cai and Clapham, 2012; Wan and Jékely, 2021). Taken together, the acquisition of Ca²⁺-regulated asymmetric wave propagation in the posterior flagellum was likely a key event in early opisthokont evolution.

## Methods

### Apusomonad culture

The apusomonad *P. kaiyoae* n. sp., originally isolated from a deep-sea sediment sample (Yabuki et al., 2023), was cultured in Hemi LB medium, supplemented with 1% v/v horse serum and 0.1% v/v LB medium in artificial seawater (Daigo’s ASW), and maintained at 20 °C. Subculturing occurred weekly in a 25 ml petri flask under a 12-hour:12-hour light:dark cycle. Cells used for experiments were ten days post-inoculation.

### Recording light-induced response of *P. kaiyoae*

The Hemi LB medium was replaced with Herbst modified Artificial Seawater (ASW) in the cell suspension. The cells were then transferred to a microchamber on a glass slide (∼1 mm depth) and kept in darkness for 1 hour to allow them to settle. ASW was composed of 462.01 mM NaCl, 9.39 mM KCl, 10.81 mM CaCl_2_, 48.27 mM MgCl_2_ and 10 mM Hepes-NaOH (pH 8.0). Cell movement was observed using an inverted phase contrast microscope (IX71, Olympus, Japan) equipped with 10× or 40× objectives. For illumination, the entire field was exposed to a laboratory-made red stroboscopic LED lamp (620-630 nm, Power LED, Edison, Taiwan), while the center area was illuminated with blue light (440 nm; Spectra X, Lumencor Inc., OR, USA) using an iris diaphragm through the objective (∼50 μmol photons m^-2^ s^-1^). In experiments involving different colors, cells were illuminated with cyan (470 nm), green (550 nm), and yellow (565 nm) light. Weak or strong blue light was used to test cell behavior at 5 μmol photons m^-2^ s^-1^ or 80 μmol photons m^-2^ s^-1^, respectively. High-speed cameras (HAS-U2, DITECT, Japan) were used to record images at 1 frame per second (fps) for trajectory analysis and 100 or 150 fps for waveform analysis. Photoavoidance was assessed by measuring the relative light intensity in the blue-light-illuminated center using ImageJ. Cell density, velocity, and flagellar waveforms were analyzed using Bohboh software (Bohboh Soft, Tokyo, Japan), while circular data analysis was performed with Oriana software (Kovach Computing Services, UK).

### Effects of drug treatment

Nocodazole (Sigma-Aldrich), ciliobrevin A (Selleck), cytochalasin B (Sigma-Aldrich), and blebbistatin (FUJIFILM Wako Pure Chemical) were prepared by dissolving in dimethyl sulfoxide to concentrations of 12.5 mM, 40 mM, 200 mM, and 10 mM, respectively. Each drug was added to artificial seawater at a 1:1000 dilution. After the cells in Herbst modified ASW settled in the microchamber, the drug-containing artificial seawater was perfused into the chamber, and cell motility was recorded 15 minutes post-perfusion.

### Effects of extracellular calcium ion concentration

Cells were loaded into a microchamber and allowed to settle. Subsequently, ASW containing one-quarter of the normal CaCl_2_ concentration (low Ca^2+^ ASW) was perfused into the chamber, and cell motility was recorded 15 minutes later. The chamber was then perfused with Herbst modified ASW, and motility was recorded again 25 minutes later to confirm cell viability and contractility.

### Antibodies and immunofluorescent microscopy

Cells were attached to glass slides and fixed with 2% paraformaldehyde for 30 minutes, followed by 4% paraformaldehyde for another 30 minutes at room temperature. After washing twice with PBS and once with PBS containing 0.05% Triton X-100 (TPBS), samples were incubated in blocking buffer (10% goat serum in TPBS) for 1 hour at room temperature. Subsequently, they were incubated with an anti-acetylated alpha-tubulin mouse antibody (T6793, Sigma-Aldrich, St. Louis, MO, USA; 1:1000 dilution) in blocking buffer for 1 hour at room temperature. Following four washes with PBS, samples were stained with a mixture of Alexa Fluor 488-conjugated goat anti-mouse antibody (Molecular Probes, Eugene, OR, USA; 1:1000 dilution) and Alexa Fluor 546 Phalloidin (A22283, Invitrogen; 1:1000 dilution) in blocking buffer for 1 hour at room temperature. In some experiments, anti-actin rabbit antibody (1854-1, Epitomics; 1:200 dilution) was used. After three 5-minute PBS washes, samples were mounted in SlowFade Gold Antifade Mountant (Thermo Fisher Scientific). Super-resolution images were obtained using structured illumination microscopy (lattice SIM) with Elyra 7 (ZEISS, Germany).

### Ca^2+^ imaging

Cells were transferred to a microchamber on a glass slide (∼1 mm depth) and kept in darkness for 2 hours. The solution was replaced with Hemi LB medium containing 0.04% Cremophor, 5 μM Fluo-8H AM, and 0.1% DMSO, followed by 30 minutes in darkness at 20 °C. After four washes with ASW, the cells were further incubated in darkness for 30 minutes. Photophobic response induction and Fluo-8H excitation were performed simultaneously using cyan light (470 nm, intensity 30 μmol photons m^-2^ s^-1^) on a microscope (IX71, Olympus) at 10 fps, using the High-Speed Recording Software (Hamamatsu Photonics). Fluorescent intensity was analyzed using Bohboh software, and pseudo-color videos were generated using ImageJ. Control experiments were conducted under the same conditions without 5 μM Fluo-8H AM.

### Electron microscopy

Thin-section electron microscopy was conducted following previously described methods (Konno et al., 2010; Sunter et al., 2019), with some modifications. Cells were harvested, centrifuged at 700 g for 5 minutes at 20°C, and fixed with 2.5% glutaraldehyde in 0.1 M cacodylate buffer (pH 7.4) for 1 hour at room temperature. After washing with 0.1 M cacodylate buffer, cells were post-fixed with 1% OsO_4_ in 0.1 M cacodylate buffer at 4 °C for 2 hours. Cells were washed five times with ddH_2_O and then stained *en bloc* with 2% aqueous uranyl acetate overnight at 4 °C. Samples were then dehydrated in a 30-100% EtOH series, substituted with propylene oxide, and embedded in Agar Low Viscosity Resin (Agar Scientific, Stansted, UK). Sections (70 nm) were cut using an ultramicrotome (LEICA, Wetzlar, Germany) and mounted on a formvar-coated grid. Samples were stained with Reynolds’ lead citrate for 5 minutes and observed using a JEM-1200EX transmission electron microscope (JEOL, Tokyo, Japan).

### Statistical analysis

Data are expressed as means ± standard deviation in box plots in R. Statistical analyses were performed using Dunnett’s multiple comparison test or Student’s T test. Statistical significance was defined as * = P < 0.05, ** = P < 0.01 or *** = P < 0.001.

### Data availability

All data needed to evaluate the conclusions in the paper are present in the paper and/or the Supplementary Materials. Additional data related to this paper may be requested from the authors.

## Acknowledgements

We thank Dr. Luis Javier Galindo, University of Oxford, for critical reading of the manuscript. This work was also supported, in part, by Grants-in-Aid for Scientific Research (A) (17H01440) and Challenging Exploratory Research (15 K14566) from the Japan Society for the Promotion of Science, Japan (JSPS); Grants-in-Aid for Innovative Areas (15H01201 and 15H01308) and Transformative Research Areas (21H05304) from the Ministry of Education, Culture, Sports, Science and Technology, Japan.

## Author contributions

K.I. designed the outline of the study. A.S, C.Y. and K.I. conducted the main experiments. Y.S. and A.Y. handled cell culture. R.Y. performed electron microscopy; R.Y., A.S., K.S. and K.I. conducted analysis. K.I. wrote the manuscript. All authors revised and approved the final version of the manuscript.

## Supplementary information

**Supplementary Figure 1**

The ausomonad *Podomonas kaiyoae* shows no positive phototaxis.

**Supplementary Figure 2**

Distributions of microtubules and actin filaments in the apusomonad *Podomonas kaiyoae*.

**Supplementary Figure 3**

Thin-sectioned electron microscopic images of *P. kaiyoae*.

**Supplementary Video 1**

Blue light induced avoidance reaction of the apusomonad *Podomonas kaiyoae*. The movie plays at a speed of 30x. Bar, 100 µm.

**Supplementary Video 2**

Illumination of green light induces less avoidance reaction of apusomonads. The movie plays at a speed of 30x. Bar, 100 µm.

**Supplementary Video 3**

Cells gliding over the boundary from illuminated bright area to the outside area. The movie plays at a speed of 6x. Bar, 100 µm.

**Supplementary Video 4**

Cells approaching from outside to illuminated area. The movie plays at a speed of 6x. Bar, 100 µm.

**Supplementary Video 5**

The apsomonad *Podomonas kaiyoae* shows no positive phototaxis. Blue light was illuminated from one side of the chamber on a glass slide (arrow). The movie plays at a speed of 30x. Bar, 100 µm.

**Supplementary Video 6**

Flagellar bending and cell contraction during the photophobic reaction of ‘settled’ apusomonads. The movie plays at a speed of 0.2x. Bar, 10 µm.

**Supplementary Video 7**

Flagellar bending and cell contraction during the photophobic reaction of ‘gliding’ apusomonads. The movie plays at a speed of 0.2x. Bar, 10 µm.

**Supplementary Video 8**

Cell contraction and posterior flagellar bending during photophobic reaction of apusomonads. The movie plays at a speed of 0.3x. Bar, 10 µm.

**Supplementary Video 9**

Response of nocodazole-treated cells to blue light. Blue light was illuminated to the center area (blue). The movie plays at a speed of 30x. Bar, 100 µm.

**Supplementary Video 10**

Response of cytochalasin-treated cells to blue light. Blue light was illuminated to the center area (blue). The movie plays at a speed of 30x. Bar, 100 µm.

**Supplementary Video 11**

Response of ciliobrevin-treated cells to blue light. Blue light was illuminated to the center area (blue). The movie plays at a speed of 30x. Bar, 100 µm.

**Supplementary Video 12**

Response of blebbistatin-treated cells to blue light. Blue light was illuminated to the center area (blue). The movie plays at a speed of 30x. Bar, 100 µm.

**Supplementary Video 13**

Adhesion of posterior flagellum on glass surface during cell flipping in cytochalasin-treated cells. The movie plays at a speed of 0.3x. Bar, 10 µm.

**Supplementary Video 14**

Ciliobrevin-induced cell flipping by blue light illumination in apusomonads. The movie plays at a speed of 0.3x. Bar, 10 µm.

**Supplementary Video 15**

Response of *P. kaiyoae* behavior to blue light in the artificial sea water containing a low concentration of Ca^2+^. The movie plays at a speed of 30x. Bar, 100 µm.

**Supplementary Video 16**

Response of *P. kaiyoae* cells to blue light in the artificial sea water containing a low concentration of Ca^2+^. The movie plays at a speed of 0.3x. Bar, 10 µm.

**Supplementary Video 17**

Changes in intracellular Ca^2+^ concentration during cell contraction. Fluo-8H fluorescence is displayed by pseudocolor. A or P indicates the anterior or posterior side of a cell, respectively. The movie plays at a speed of 0.16x. Bar, 10 µm.

