## Supplementary Figures for "Apusomonad photophobic behavior highlights cytoskeletal responses to blue light in early eukaryotes"

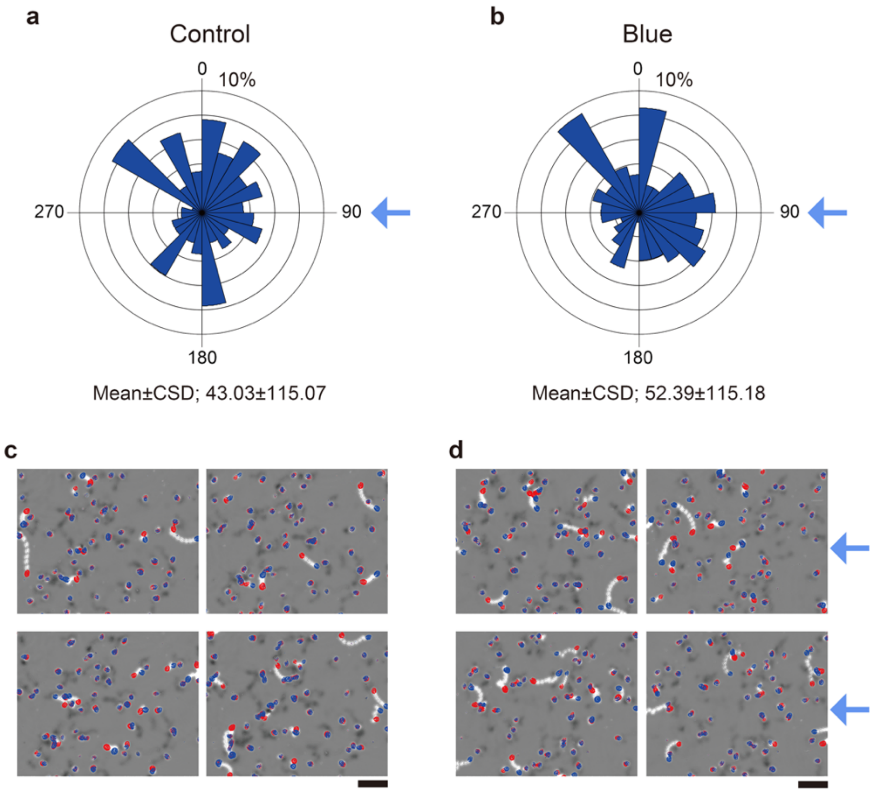


**Supplementary Figure 1**

The ausomonad *Podomonas kaiyoae* shows no positive phototaxis. **a, b,** Rose plots showing the gliding direction under stroboscopic illumination of red light without (a, control, n=128) or with (b, blue, n=118) blue light from one side of a chamber on a glass slide. CSD, circular standard deviation. Arrow shows the direction of blue light. **c, d,** Trajectories of gliding cells under illumination of red light without (c) or with (d) blue light from one side of a chamber on a glass slide. Ten images acquired at 1 s intervals are superimposed. Red and blue dots indicate the start and end points of the superimposition, respectively. Arrows show the directions of blue light. Scale bar, 100 μm.

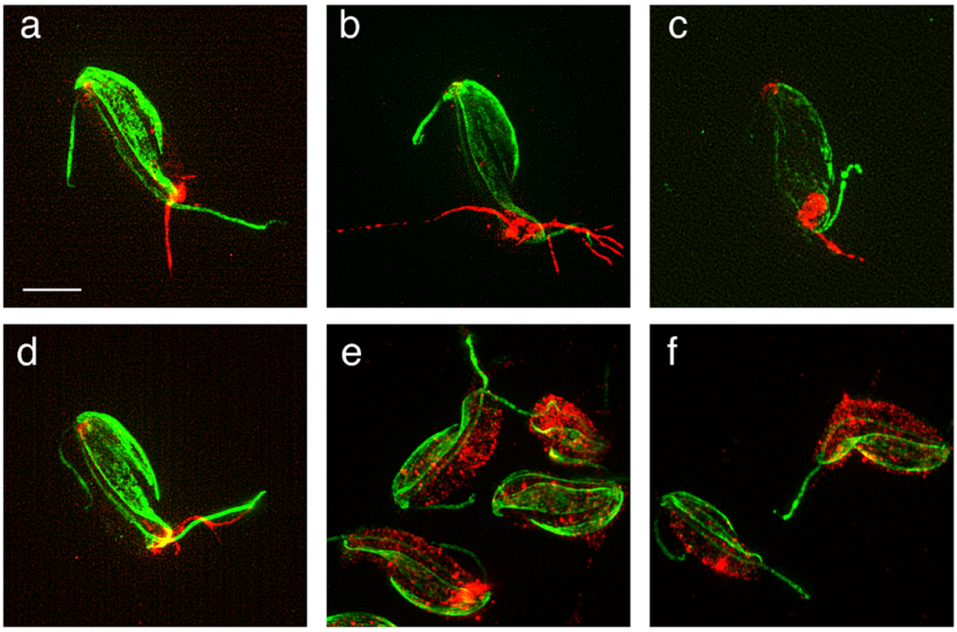


**Supplementary Figure 2**

Distributions of microtubules and actin filaments in the apusomonad *Podomonas kaiyoae*. Cytoskeletons were visualized by immunofluorescence staining with anti-acetylated α-tubulin antibody (green) and Alexa Fluor 546 phalloidin (red) (a-d), and with anti-acetylated α-tubulin antibody (green) and anti-actin antibody (e-f). Image a is the same as Fig. 3a. Bar, 5 µm.

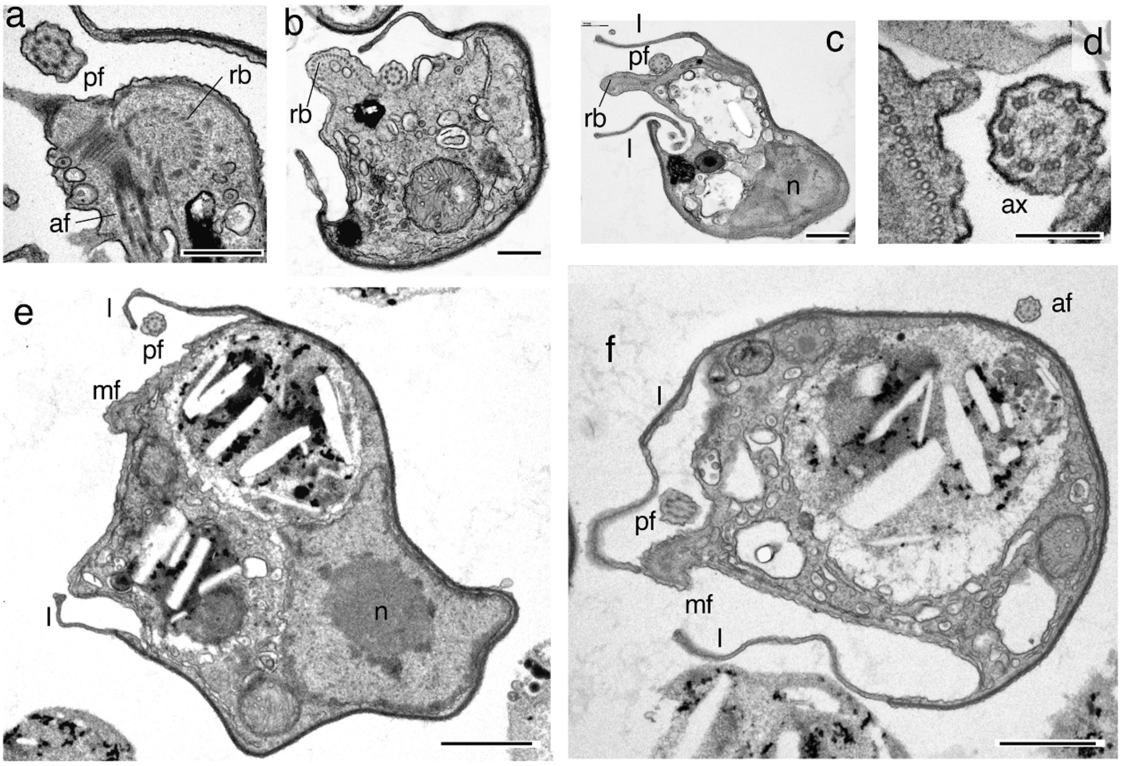


**Supplementary Figure 3**

Thin-sectioned electron microscopic images of *P. kaiyoae*. af, anterior flagellum. ax, axoneme. l, lip. mf, microfilament bundle. n, nucleus. pf, posterior flagellum. rb, ribbon. Bar, a-c: 500 nm; d, 200 nm; e-f, 1 μm.
